# PROPER UTILIZATION OF IODIZED SALT AND ASSOCIATED FACTORS AMONG RURAL COMMUNITY OF HETOSA DISTRICT, OROMIA REGIONAL STATE, SOUTH EAST ETHIOPIA

**DOI:** 10.1101/2020.06.04.133926

**Authors:** Abu Tura Bulli, MeleseTadese Aredo, Hailu Fekadu, Ashenafi Habtamu Regesu

**Author notes:** Corresponding Authur:Abu Tura Bulli.

## Abstract

**Introduction:** Iodine is considered to be one of the most essential micronutrients for the normal physical and mental development of human beings. However, little is known about households’ use of iodized salt and associated factors.

**Objectives:** This study was to assess the proper utilization of iodized salt at the household level and associated factors in Hetosa District, Southeast Ethiopia, 2019.

**Methods:** A Community-based cross-sectional study was conducted from August 20 up to September 15/2019 in rural Hetosa District, Arsi Zone, and east-south Ethiopia. A total of 603 households were selected using a systematic random sampling technique. Data were collected employing structured and pre-tested questionnaires by face -to -face interview technique. The use of iodized salt at the household level was tested with the iodine rapid test kit. The data were checked, coded and entered into Epi Info Version 7 and export to SPSS version 21 for analysis.

**Result:** A total of 596 participants were included in this study. The availability of adequately iodized salt was 61.1%. The proportion of proper utilization of iodized salt at the household level was 38.4%.Formal Educational (AOR=1.688, 95%CI (1.002, 2.846)),Practice of iodized salt use (AOR= 3.352, 95%CI (2.160, 5.202)), Knowledge on use of iodized salt (AOR=2.320, 95%CI (1.437, 3.745)) and level of iodine content in salt (AOR= 1.668, 95%CI (1.071, 2.597)) were statistically significant to utilization of iodized salt.

**Conclusion:** Proper utilization of iodized salt remains very low, which was 38.4% in the district and does not meet the national goal. Educational status, level of iodized salt, good knowledge and good practice were significantly associated factors with proper utilization of adequately iodized salt in this study.

## 1. Background

Iodine is an essential micronutrient. The human demand for iodine is a hundred and fifty mcg/ day.Iodine is a trace element sparsely distributed over the surface of the earth. About 90% of this comes from food while 10% from water. It is essential for the synthesis of the thyroid hormone, which is necessary for human growth and development***(1)***. Iodine is present in the body in a minute amount, mainly in the thyroid gland. Its confirmed role is in the synthesis of thyroid hormone(2).

Iodine is an essential dietary nutrient that helps the body to manufacture thyroxin, the hormone that regulates normal growth and development. The quantity of iodine required by an individual is minute, being 150–200μg per day or a teaspoonful during a lifetime(4). Iodine is considered to be one of the most vital micronutrients for the standard physical and mental changing of anthropological existences. The erosion of soil and deforestation is considered to be the reasons for the decrease of the iodine level in hilly and mountainous areas(5).Universal salt iodization (USI) is globally accepted as the most cost-effective public health strategy to prevent iodine deficiency(6).This strengthening salt iodization programs and improving monitoring is a crucial step to eradicate the problem(7).

Many studies evidenced that iodine deficiency disorders contribute to the increment of morbidity and mortality of infants and neonates. It also reduces the quality of life, national productivity and 13.5 points of the intelligent quotient (IQ)(22). If iodine deficiency continued as a problem by 2012-2025 about 12 million children would be born from iodine-deficient mothers; these children would have some point of permanent brain damage (with a decrease in IQ). Economic productivity losses related to iodine deficiency during 2012-2025 is US$5 billion (5).

The prevalence of IDD based on the total goiter rates (visible and palpate) is the highest in the Eastern Mediterranean region 32%, followed by Africa 20%, European 15% and Southeast Asia 12%. However, it is important to recognize that these clinical signs are superficially in severe cases and subclinical deficiency, which is also associated with a range of intellectual and behavioral deficits, affects many more individuals. Nearly one -third of the world’s population (2.3 billion is considered to be at risk of iodine deficiency, which a large proportion live in Southeast Asia(22).

According to World Health Organization (WHO),United Nation International Child Fund(UNICEF) and International Council for Control of Iodine Deficiency Disorder (ICCIDD) the daily recommended dietary allowance (RDA) is 90, 120, 150, 200 mcg of iodine for preschool children, school children, adults, and pregnant and lactating women respectively(10).

Ethiopian Public Health Institute 2016 survey reported that national iodized salt utilization coverage was 89.2%.However,only about 26% of the surveyed households had salt that was adequately iodized salt (at≥15ppm)(13).The National goal of iodization utilization coverage of Ethiopia 95%(14).

It is estimated that almost half of Ethiopia’s 80 million population faces iodine deficiency disorder (IDD), raising alarm in the Horn of Africa nation(23).In Ethiopia, one out of each a thousand may be a hypothyroidism and concerning 50,000 prenatal deaths were occurring annually because of iodine deficiency disorders, twenty sixth of the overall population have a goiter and 62% of the population is at risk of IDD according to the national survey made by the previous Ethiopian Nutrition Institute (24).

Study conducted in Chole district arsi zone south east Ethiopia,among school children revealed that the prevalence of goiter was 36.6%(25).A study conducted on the Utilization of Iodized Salt at Household Level in Zuway Dugda district revealed that, only 25.6% nof the respondents properly utilized iodized salt(26).

Therefore,eventhough iodine deficiency is the major public health problems and the cause of morbidity and mortality, but very little is known about households’ use of iodized salt and associated factors in Ethiopian context and why this study was conducted.

## 2. Methods and Materials

### 2.1. Study area

The study was carried out in Hetosa district one of the 26 districts found in Arsi Zone. Hetosa district is located in Arsi zone in Oromia Regional State, about 160 Kilometers Southeast from Addis Ababa of Ethiopia. The area lies between 08° 08’N-08°13’ E latitude and 39° 14’ N -39° 23’ E longitude with an elevation range from1500-4170meters above sea level and characterized by mid subtropical temperature ranging from 5 C°-28 C°. According to the District Health Office 2019 report, there are four Primary Health Care Unit which includes four health centers and 23 health posts. The potential health coverage by health center 62.7% in 2019(55).

### 2.2. Study design and period

The community-based cross-section study design was carried out at household level among a rural community of Hetosa District, Southeast Ethiopia. The study was carried out from August 20 up to September 15/2019.

### 2.3. Source and *Study* population

#### 2.3.1 Source population

The source populations were all households who live in Hetosa District

#### 2.3.2 Study population

Households were systematically selected in the selected Kebele of the Hetosa District to be part of the study.

**Inclusion criteria:** The study was conducted among adult aged 18 years and above those who involved in food item purchasing and preparation/cooking and live the area for more than 6 months.

**Exclusion Criteria:** The household head/food preparation had seriously sick that makes communication difficult to get the necessary information

### 2.4. Sampling procedure

In this study multi-stage sampling method was used, by considering Hetosa district, which consists of twenty-three Kebele. Kebele were selected using simple random sampling from the total 23 rural administrative areas (Kebele) of the district, from those Kebele 30% (7 Kebele).The calculated sample size (603 households) were proportionately allotted to every elect Kebele supported its size of households. Then a systematic random sampling technique was used to identify the study households from each selected Kebele.

The first house was chosen at randomly by lottery method. This house became the starting point or first house to sampling frame. Where there was more than one eligible individual in selected households, a lottery method was used to pick one of them. Where there were 4 or more eligible households in a house selected, two households was drawn at random and only one eligible person (household head) a lottery method was used to pick one of them. Where no eligible household was found in a selected house, the next house closest to the previously selected one was selected. This procedure was repeated until the required number of respondents was obtained. When the interviewer refused to involve in the study, the household was taken as non-response.

### 2.5. Variables of the study

#### 2.5.1 Dependent variable

Proper utilization of iodized salt

#### 2.5.2 Independent variables

**Socio-demographic factors**: Sex, Responsibility in the Household, Ethnicity, Religion, Age, Educational status, Family income, Family size, marital status, Occupational status, House type

**Behavioral factor**:Knowledge about iodized salt,Practice in about the iodized salt in the community,Time salt to add at cooking,Source of information

**Environmental factor**: Place of Storage salt, expose to sunlight, Duration of storage, Salt container cover, place of usually buy iodized salt, Type of salt used,Time travel to buy salt

**Health and health-related system**: Availability of iodized salt

### 2.6. Data collection procedures (instruments, personnel, measurements)

Data were collected using a structured questionnaire and rapid iodine test kit by face-to-face interview with four diploma nurse and Supervised by 2 BSC Nurse. The standard structured questionnaire which was modified for this study was used for recording the responses by the interviewee. The questionnaire was adapted in English and then translated to Afan Oromo and finally, it was retranslated in to English by another language translator for checking consistency. The questionnaires were used to collect data on socio-economic characteristics, Behavioral factors, and environmental factors associated with utilization of iodized salt.

For the data collector training was given for one day, focusing on communication and technical issues on how to collect data, interview technique and recording how to test the salt for adequately iodized and supervisors was trained on how to supervise the data collectors by the investigator. Questions was analyzed and reported before actual data collection on similar kebelesbut not included in the study subject for completeness, consistency, and timeliness. Data collection was started with a brief introduction and after consent for participants obtained and then questionnaire completed by face to face interview, in the language of respondent and salt tested.Supervisors were checking questionnaire and contact at least 50% of the respondents to verify that the correct procedures had been followed in data collection and testing for iodized salt.

Salt test kit –To assess the availability of adequately iodized salt at household, the interviewer asks every sample household to provide a teaspoon of salt used for food preparation and fill small cup spread flat, add two drop of test solution on surface of salt by piercing the white ampule and compare color on salt with color chart with one minute and determine the iodine concentration.

If no color change appeared on the salt (after one minute, on a fresh sample add up to 5 drop of recheck solution in the red ampule and the 2 drops of test solution on the same spot and compare the color with color chart and determine iodine content. The sample sale was taken from the salt container and tested for its adequate iodine concentration, intense color. This determined by using improving iodized salt field test kit.For salt fortified with potassium iodate only and range from no iodized salt (0 ppm) inadequately iodized salt(less than 15ppm)light blue, and adequate iodized salt (greaterthan15ppm) deep blue. The unopened ampule was used and the kit was accurate for visual detection of potassium iodate concentration at the threshold of 15ppm and result was valid. The test kit was obtained from Arsi Zonal Health Department; Rapid test kit one vial can test approximately 50 salt sample table, bags or packages for the salt sample.

### 2.7. Data quality assurance

Questionnaire was checked and screened for completeness and consistency. Omission, error, completeness was checked. Data were cleaned, edited and coded before entry. Data processing was done before analysis. The prepared questionnaire was pretested prior to the actual data collection on 30 participants. Checking on the spot and double data entry on EPI INFO-7 was done to ensure the completeness and consistency of the information collected.

### 2.8. Data processing and analysis

Data were entered, cleaned and edited by using EPI-INFO Version -7 Statistical software and then data was exported to SPSS version 21 for analysis. Descriptive statistics of the collected data was done for most variables in the study using statistical measurements. Frequency table, graph, percentages, means, and standard deviation were used. Bivariate & multivariate analyses were performed using binary logistic regressions, to identify the relationship between dependent & independent variables. Candidate variables with P-value **<**0.25 in bivariate analysis and those considered important based on literatures were entered into multivariate regressions. The final model was built with backwards selection & a significant association was declared at a p-value less than 0.05. Finally, the results were presented in texts and tables with corresponding AOR, 95% CI & P-value.

### 2.9. Ethical considerations

Ethical clearance was obtained from the ethical review board of Arsi University and the formal letter of support obtained from the Hetosa Woreda. Verbal consent was obtained from households head and study participants. The participants were allowed to consider their participation and given the opportunity to withdraw from the study at any point in the course of study when they were not comfortable with the study. Data collector presented questions for the respondent by their language. Name of the respondent was not including in the questionnaire and information of individual respondent was not being shared to ensure confidentiality. All the responses of the participants and the results obtained from each household were kept anonymous and confidential by using a coding system whereby no one should had access except the principal investigator. The study had no risk on the study subject and interview was conducted in a private area.

### 3. Result

### 3.1. Socio demographic characteristics of respondent’s

From 603 participants invited to participate in the study, 596 participants were included with a response rate of 98.8%. The mean age ± standard deviation participants were 34 ± 11.8 years. The age of participants ranged from 18 to 60 years. Almost 577(96.8%) were female, 585(98.2%) participants were Oromo by ethnicity group, 382(64.1%) were Muslim by religion, 508(85.2%) were Married, 498(83.6%) were housewife and 281(47.1%) had at least Primary Education (1-8) level, 351(58.9%)participants had less than 5 family size,209(35.1%) of participants monthly income was greater than 1500 ETB and 508(85.2%) participants had the iron sheet house(Table 1).

**Table 1:**
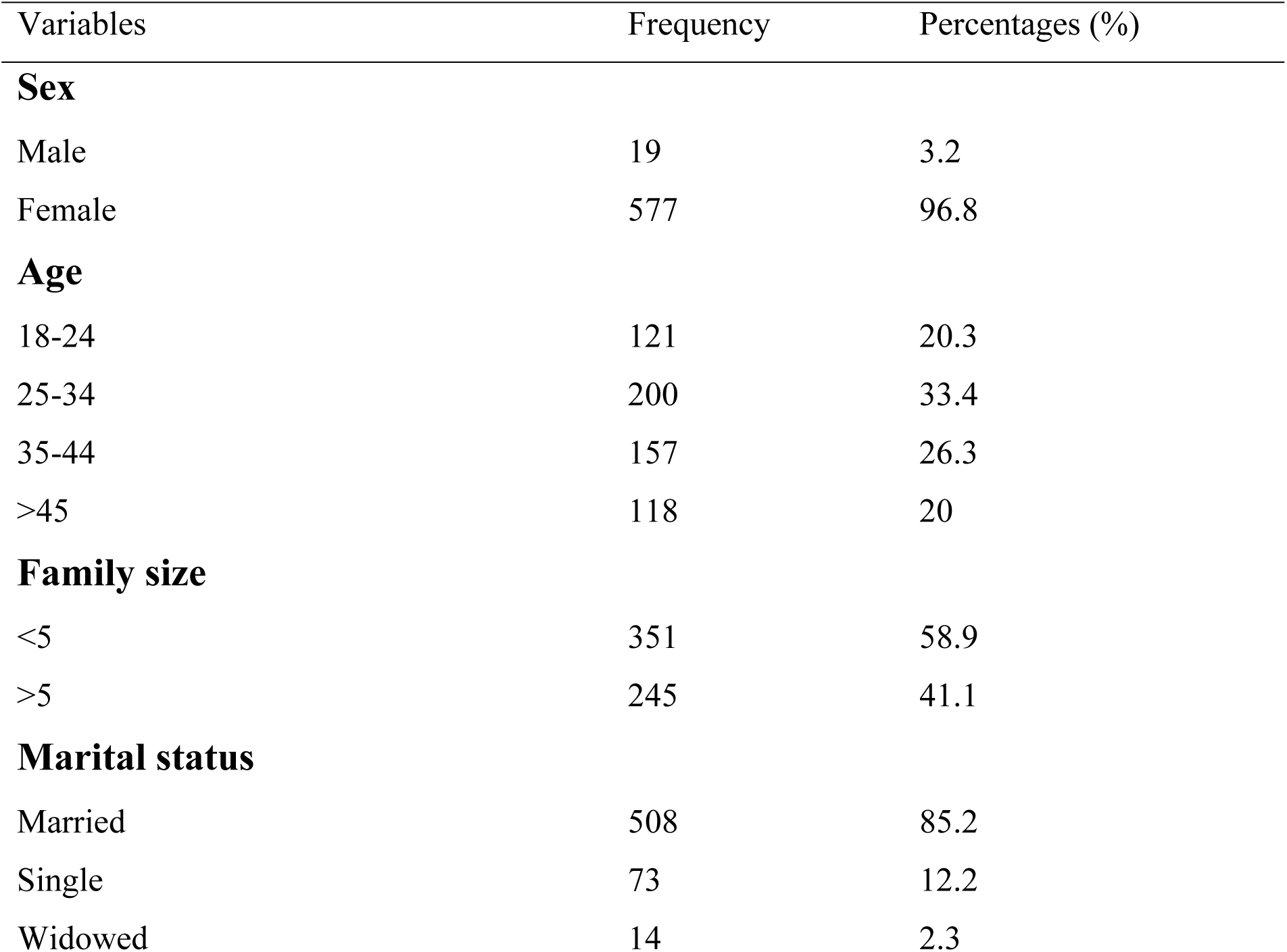
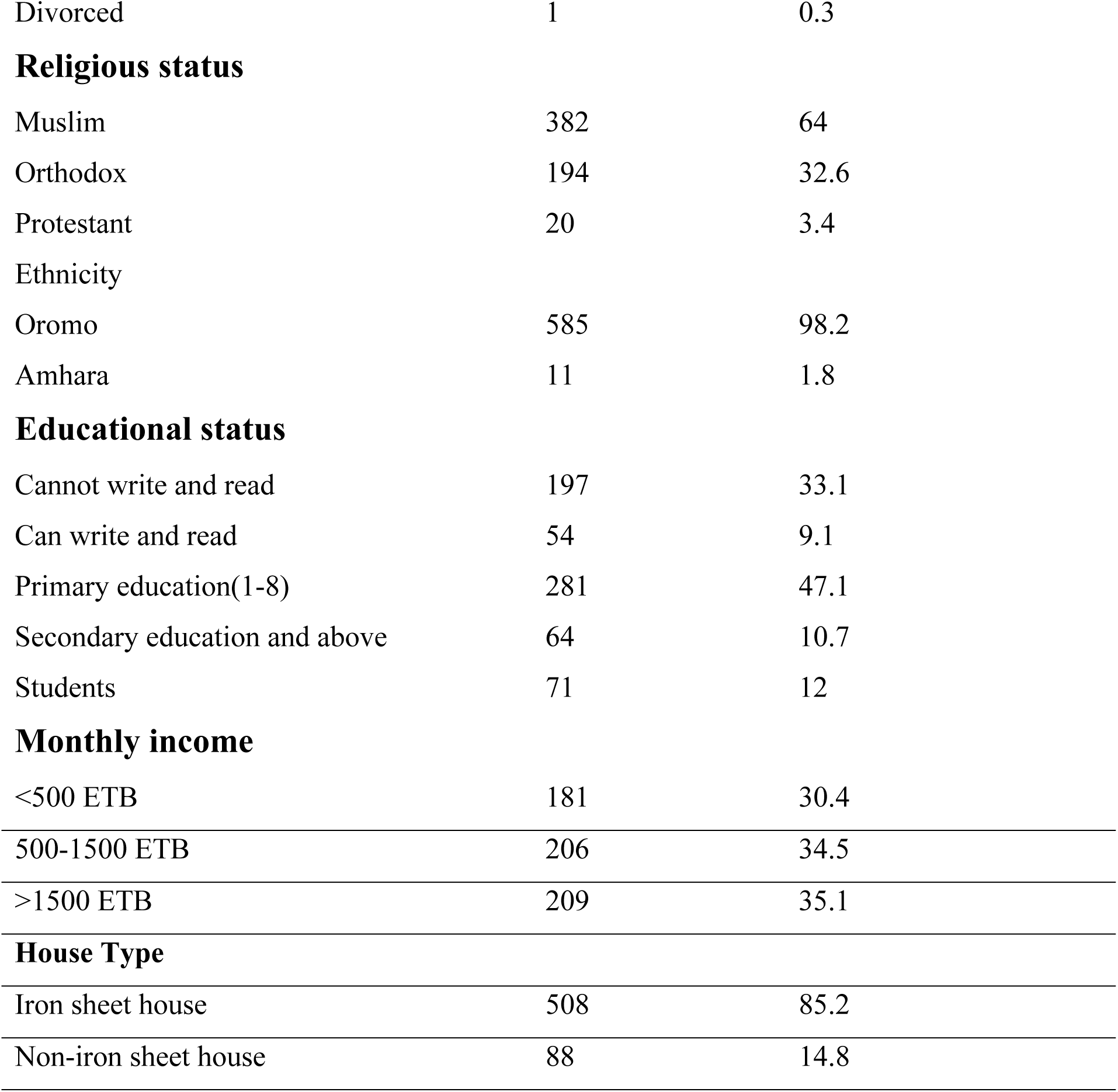
Descriptive analysis of Socio demographic and economics characteristics of respondent’s utilization of iodized salt at households and associated factors in Hetosa District, Southeast Ethiopia, August 2019(n=596)

| Variables | Frequency | Percentages (%) |
| --- | --- | --- |
| **Sex** |  |  |
| Male | 19 | 3.2 |
| Female | 577 | 96.8 |
| **Age** |  |  |
| 18-24 | 121 | 20.3 |
| 25-34 | 200 | 33.4 |
| 35-44 | 157 | 26.3 |
| >45 | 118 | 20 |
| **Family size** |  |  |
| <5 | 351 | 58.9 |
| >5 | 245 | 41.1 |
| **Marital status** |  |  |
| Married | 508 | 85.2 |
| Single | 73 | 12.2 |
| Widowed | 14 | 2.3 |
| Divorced | 1 | 0.3 |
| **Religious status** |  |  |
| Muslim | 382 | 64 |
| Orthodox | 194 | 32.6 |
| Protestant | 20 | 3.4 |
| **Ethnicity** |  |  |
| Oromo | 585 | 98.2 |
| Amhara | 11 | 1.8 |
| **Educational status** |  |  |
| Cannot write and read | 197 | 33.1 |
| Can write and read | 54 | 9.1 |
| Primary education(1-8) | 281 | 47.1 |
| Secondary education and above | 64 | 10.7 |
| Students | 71 | 12 |
| **Monthly income** |  |  |
| <500 ETB | 181 | 30.4 |
| 500-1500 ETB | 206 | 34.5 |
| >1500 ETB | 209 | 35.1 |
| **House Type** |  |  |
| Iron sheet house | 508 | 85.2 |
| Non-iron sheet house | 88 | 14.8 |

### 3.2. Knowledge on the Proper utilization of iodized salt at the household level

From the respondents, 382(64.1%) heard about the iodized salt. In this study from the total respondents nearly half of the 374(62.8%) not heard about Iodine Deficiency Disorder. Among the respondents who heard about Iodine Deficiency Disorder 69(12%), their source information is Radio. Among the respondents, only 236(39.6%) knew the benefit of iodized salt. Out of those respondents, the question only 31.5% knew Goiter result from iodine deficiency and 30% knows that regular consumption of iodized salt prevents iodine deficiency in the body.

Among those interviewed participants most of them 457(76.7%) didn’t know what Problem occurred when not consumed Iodized salt at the household level. From the total respondents, 182(30.5%) had good knowledge about the proper utilization of iodized salt at the household level (Table 2).

**Table 2:** Knowledge of respondent’s regarding the importance of utilization of iodized salt inrural communities of Hetosa District, Southeast Ethiopia, August 2019(n=596).

| Variables | Frequency | Percentages (%) |
| --- | --- | --- |
| **Have heard about iodized salt** |  |  |
| Yes | 382 | 64.1 |
| No | 214 | 35.9 |
| **Have heard about IDD** |  |  |
| Yes | 222 | 37.2 |
| No | 374 | 62.8 |
| **Source of information on IDD** |  |  |
| TV | 60 | 10.1 |
| Radio | 69 | 12 |
| HEW | 54 | 9.1 |
| Magazine | 16 | 2.7 |
| From Family | 23 | 3.9 |
| **Knowledge on the benefit of iodized salt** |  |  |
| Yes | 236 | 39.6 |
| No | 360 | 60.4 |
| **Goiter results from iodine deficiency** |  |  |
| Yes | 188 | 31.5 |
| No | 408 | 68.5 |
| **Regular Consumption of iodized salt prevent IDD in the body** |  |  |
| Yes | 179 | 30 |
| No | 419 | 70 |
| **Problem occurred when not consumed Iodized salt** |  |  |
| Mental retardation | 126 | 21.1 |
| Abortion | 4 | 0.7 |
| Infant mortality | 9 | 1.5 |
| Unknown | 457 | 76.7 |
| **Knowledge on IDD and use of iodized salt** |  |  |
| Poor | 414 | 69.5 |
| Good | 182 | 30.5 |

### 3.3. Utilization of iodized salt at household level

In this study among the total respondents 375(62.9%) of them, bought iodized salt in their home during the study period. Among those households which had iodized salt 211(35.4%) of the users usually buy it from the open market. From those bought the iodized salt 78.5% of the respondents time travel to buy iodized salt greater than or equal to 60 minutes. Around the 55.2% of the respondents were using non-packed salt and 44.8% of the interviewed respondents were using the packed salt. Concerning the duration of storage practice, 95% stored their salt less than 2months. Almost all 97.5% of the respondents stored salt in a dry place and 92.1% of HHs store salt in a covered container. A majority of the respondents 95.5% not exposed salt to sunlight. In this study, the respondents add salt at the end of cooking 38.4% (Table 3).

**Table 3:** Utilization of iodized salt at households level and associated factors in Hetosa District, Southeast Ethiopia, August 2019(n=596)

| Variables | Frequency | Percentages (%) |
| --- | --- | --- |
| **Do you buy iodized salt in your home** |  |  |
| Yes | 375 | 62.9 |
| No | 221 | 37.1 |
| **Place usually buy iodized salt** |  |  |
| Retail shop | 97 | 26 |
| Open market | 211 | 56.3 |
| Store | 67 | 17.7 |
| **How often you buy iodized salt** |  |  |
| Not use | 2 | 0.005 |
| Always | 111 | 30 |
| Sometimes | 262 | 69.995 |
| **Time travel to buy iodized salt** |  |  |
| <60 minutes | 128 | 21.5 |
| ≥60 minutes | 468 | 78.5 |
| **Perceived cost** |  |  |
| Expensive | 219 | 36.7 |
| Cheap | 377 | 63.3 |
| **Type of salt used in the home** |  |  |
| Packed | 267 | 44.8 |
| Non packed | 329 | 55.2 |
| Salt type most of time used in Home |  |  |
| Non-iodized salt | 267 | 44.8 |
| Iodized salt | 147 | 24.7 |
| Both | 182 | 30.5 |
| **Duration of salt stored** |  |  |
| <2 month | 566 | 95 |
| ≥2month | 30 | 5 |
| **Storage place** |  |  |
| Dry place | 581 | 97.5 |
| Moisture area | 15 | 2.5 |
| **Exposed to sunlight** |  |  |
| Yes | 27 | 4.5 |
| No | 569 | 95.5 |
| **Stored with covered Container** |  |  |
| Yes | 549 | 92.1 |
| No | 47 | 7.9 |
| **Time salt added to cooking within the last twenty four hour** |  |  |
| Early at the beginning of cooking | 367 | 61.6 |
| At the end of cooking | 229 | 38.4 |

### 3.4. Proportion of adequately iodized salt at household’s level

Salt samples were collected from 596(98.8%) participants’ households. The result of iodized salt Field Test Kit showed that from the total of 596 participants, who were requested to bring a sample of their salt, more than half of 364(61.1%) of salt samples had adequate iodine content(≥15ppm) and 17.9% of the salt sample had zero iodine contents. Out of the salt sample, 21% of the salt sample had inadequate iodine contents (<15ppm) (figure 1).

**Figure 1:**
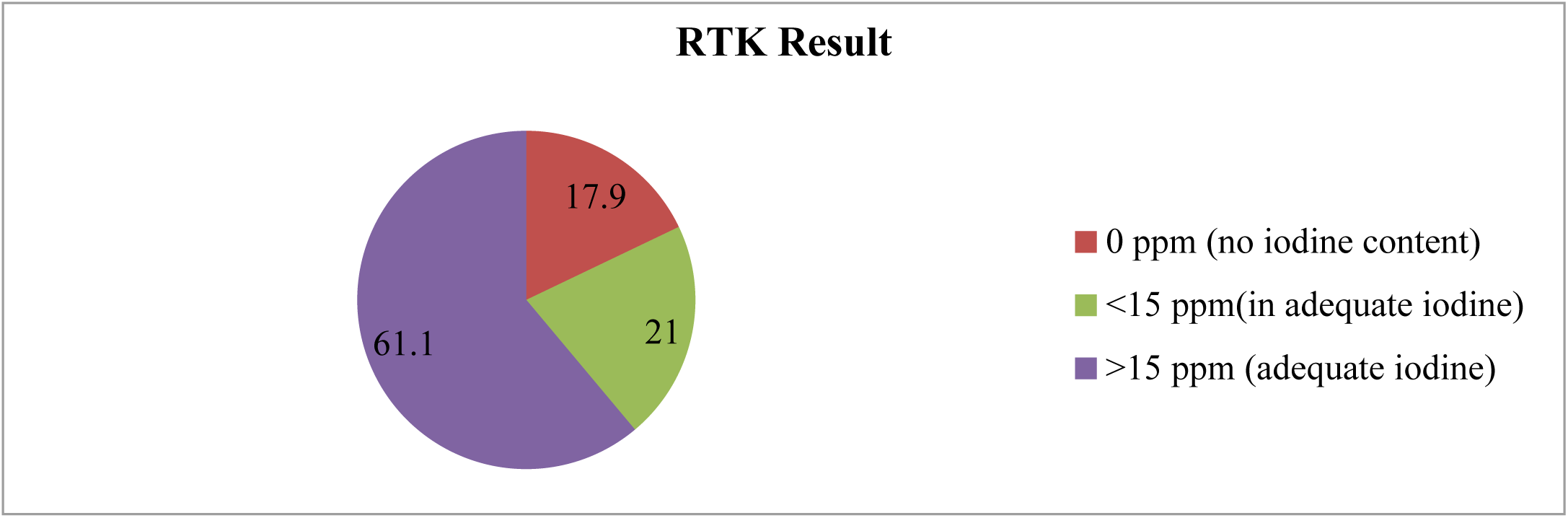
Iodine content test result by RTK at household level of communities in Hetosa District (n=596)

### 3.5. Factors Associated with Proper Utilization of Iodized Salt at Household Level

In a bivariable analysis, Age, family size, marital status, educational status, religion status, monthly income, type of house Iodine content level, knowledge, and practice were significantly associated with proper utilization of iodized salt selected as a candidate predictors for the multivariable model at P≤0.25.

In multivariable analysis after controlling the possible confounders; among variable studied, in the bivariable analysis of the utilization of Iodized salt in this study, some variables show significant differences with a p-value less than 0.05. To avoid missing variables those were significant in another similar study p-value less than 0.05 was used as a cut point off for multivariable analysis 10 variables that were significant at p-value less than 0.25 were entered into entering Logistic regression which controlled undesirable effect of confounder between variables. Hosmer and Lemeshow goodness of fit statistics were checked for the fulfilled of the model. The final model showed 0.624goodness. Vary from 0.05 and indicating the outcome variables were fully explained by the independent variables in the full model. From entered variables, only 4 variables remained a significant predictor of the outcome variables. During multivariable analysis, compared to those who did not have formal education, those who had formal education were 1.688 times more likely to use iodized salt and statistically significant compared to their counterparts[AOR=1.688(1.002, 2.846)]. The adequate level of iodized salt at household level was significantly increased. The odds of proper utilization of iodized salt by 66.8% compared to those households who had an inadequate level of iodized salt compared to their counterparts [AOR= 1.668(1.071, 2.597)]. Proper utilization of iodized salt was about two and four-folds significantly higher among those who had good knowledge compared to their counterparts [AOR=2.320(1.437, 3.745)] and good practice of iodized salt compared to their counterparts [AOR= 3.352(2.160, 5.202)] respectively (Table 4).

**Table 4:**
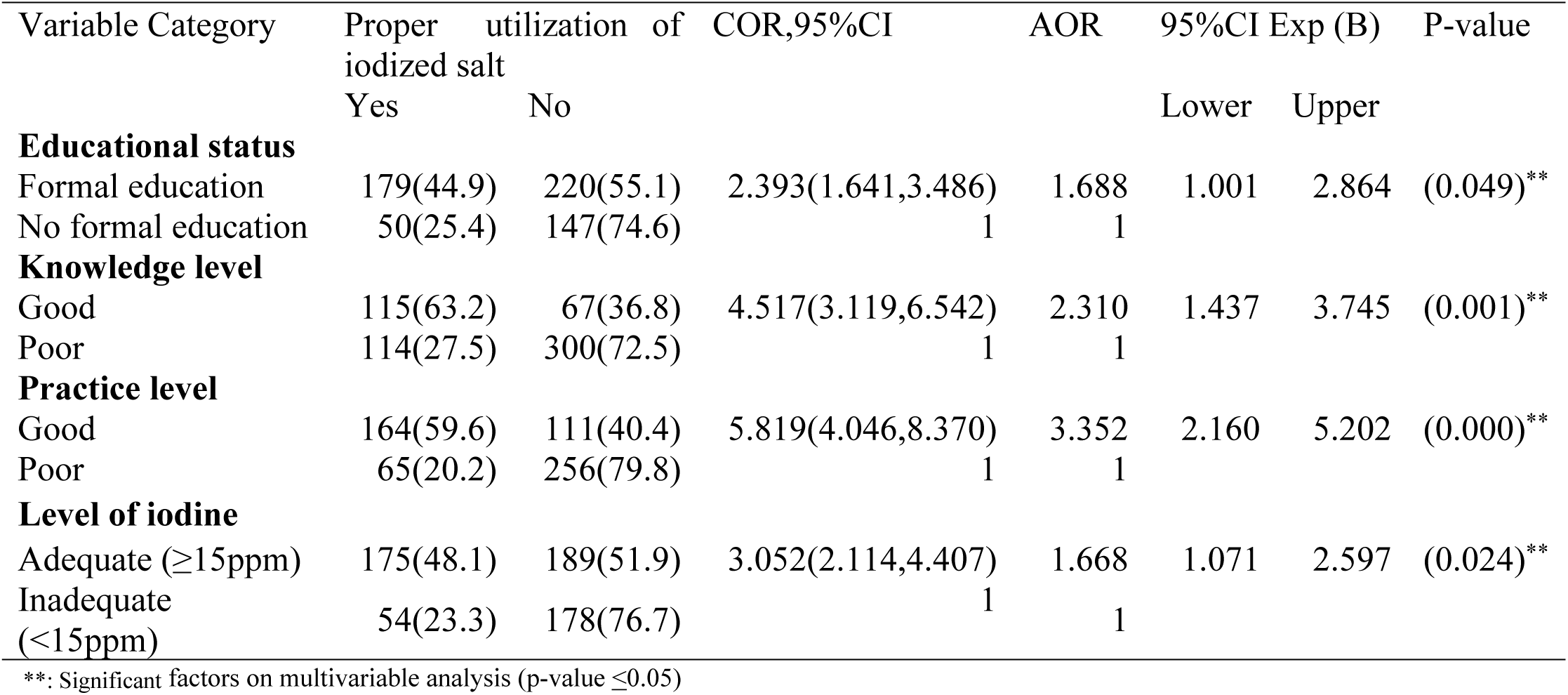
Final model of predictors of Utilization of Iodized Salt in multivariable logistic regression in Hetosa District, Southeast Ethiopia, August 2019(n=596)

| Variable Category | Proper utilization of iodized salt |  | COR,95%CI | AOR | 95%CI Exp (B) |  | P-value |
| --- | --- | --- | --- | --- | --- | --- | --- |
|  | Yes | No |  |  | Lower | Upper |  |
| **Educational status** |  |  |  |  |  |  |  |
| Formal education | 179(44.9) | 220(55.1) | 2.393(1.641,3.486) | 1.688 | 1.001 | 2.864 | (0.049)** |
| No formal education | 50(25.4) | 147(74.6) | 1 | 1 |  |  |  |
| **Knowledge level** |  |  |  |  |  |  |  |
| Good | 115(63.2) | 67(36.8) | 4.517(3.119,6.542) | 2.310 | 1.437 | 3.745 | (0.001)** |
| Poor | 114(27.5) | 300(72.5) | 1 | 1 |  |  |  |
| **Practice level** |  |  |  |  |  |  |  |
| Good | 164(59.6) | 111(40.4) | 5.819(4.046,8.370) | 3.352 | 2.160 | 5.202 | (0.000)** |
| Poor | 65(20.2) | 256(79.8) | 1 | 1 |  |  |  |
| **Level of iodine** |  |  |  |  |  |  |  |
| Adequate (≥15ppm) | 175(48.1) | 189(51.9) | 3.052(2.114,4.407) | 1.668 | 1.071 | 2.597 | (0.024)** |
| Inadequate (<15ppm) | 54(23.3) | 178(76.7) | 1 | 1 |  |  |  |

\*\* : Significant factors on multivariable analysis (p-value $\leq 0.05$ )

## 4. Discussion

Proper utilization of iodized salt at household level is a critical point to achieve the intended intervention of iodine Fortification. Universal salt iodization (USI) is globally accepted as the most cost-effective public health strategy to prevent iodine deficiency(6). This strengthening salt iodization programs and improving monitoring is a crucial step to eradicate the problem(7). Accordingly, the present study identified the overall level of proper Iodine utilization and factors associated with this proper utilization of iodine; namely, Educational status, level of iodine in salt, good knowledge, and good practices. Identifying the proper utilization of iodized salt was the final target to tackle IDDs and this means that iodized salt should be added after cooking finished in community and at households’ level (14).

Consequently, in the present study, only 38.4% of the households in the Hetosa District used a proportion of proper utilization of iodized salt. This finding was higher than the study conducted in the Province of Kenya which was 22.6% in 2015(33).The discrepancy might be due to the study period. Another study conducted in Mecha District 25.7% in 2019, Ahferom district 8.9% in 2012, Sideman zone Bensa Woreda 10.6% in 2016, Zuway Dugda district 25.7% in 2019(14, 26, 38, 46)) of them utilized properly, which was lower than this study area. The difference could be due to knowledge status in the case of the Mecha District. The difference could be due to the study technique in the case of the Ahferom district. The difference could be due to the study period and design in the case of Sideman zone Bensa Woreda. The discrepancy might be due to in geographical location, educational status of the respondents, inaccessibility of awareness creation and health promotion in the case of Zuway Dugda district.

In contrary our finding is lower than study done in Andhra Pradesh Indian 48% in 2019,Laelay May chew district (59.7%) in 2015, Asella town 76.8% in 2016,Goba town 57.2% in 2016, Dega Damot 88.8% in 2019(29, 37, 40, 41, 45). This difference might be due to the study period, study area, information access, Knowledge of about iodized salt of respondents. This finding also Which is far lower than the Ethiopian Public Health Institute 2016 survey reported that national iodized salt utilization coverage was 89.2%((13).

Yet the current rate was considerably below the national goal of 95% of coverage(14).This discrepancy due to this study focus on local than the national level, the small size of area and study period. Promotions on the media increase public awareness and alert that all salt producers and traders duly iodize their salt which is essential for achieving the USI goal(7, 41). Therefore observed proper utilization of iodized salt in the study area was very low.

World Health Organization (WHO) recommended that at least 90% of households using adequately iodized salt which contains 15ppm or more at the household’s level(11). Elimination of Iodide deficiency is possible if more than 90% of the households consume adequately iodized salt and must granted(33).Consequently, the current study pointed that 61.1% tested salt have more than 15ppm iodine contents. This finding was higher compared to the study conducted in Parkasan, Andhra Pradesha Indian (42%) in 2019, Kenya (26.2%) in 2015, LaeylaMaychew (33%) in 2015, Lalo Asab (8.7%) in 2016, Afherehom district(17.5%) in 2012,(29, 33, 40, 46, 47) the households sampled in the Hetosa District had access to adequately iodized salt. The discrepancy due to income of the community, Educational status, geographical location, study period and study technique. This finding result which was lower compared to study done in Iraq (68.3%) in 2012, Asella (62.9%) in 2016, and Dessie (68.8%)in 2018(32, 41, 44).This discrepancy might be due study period in case of Asella and Dessie.

In addition the discrepancy might be due Access of information, availability packed salt, study design and residence of the respondents(37, 43). This finding, Which is far from the WHO recommendation to eliminate Iodine Deficiency Disorder 90%(10) and with the Ethiopian National Plan to eliminate IDD virtually by the year 2013 through adequate Iodization Universal Salt 90%(45). The discrepancy might be due to in educational status of respondents, Availability of the adequately iodized salt.

Level Education is one the determinant factors which facilitates or hinder the proper utilization iodized salt in community(33). Once the iodization of salt for human use becomes mandatory, periodic public education for proper storage and usage of iodized salt should continue. Public awareness and publicity is also necessary so that there is a demand for iodized salt (17).Accordingly, this study identified that the study participants who attended formal education were 1.688 times more likely to use iodized salt than those who didn’t attend informal education [AOR=1.688(1.002, 2.846)].

This finding was also similar with the study done in Sudan in 2017, Iraq in 2012, Laelay Maychew in 2015, Woilata in 2018, Asella in 2016(17, 27, 32, 40, 41), which identified education as predictor’s variables to utilization of iodized salt. On the other hand, the finding of this study odds lower than odds that studied in Province of Kenya (3.22) in 2015, Debra tabor (2.28) in 2019, Mecha district (2.2) in 209, Sideman zone (3.34) in 2016(14, 33, 38, 52).

This might be due accessibility of information, on the other hand workload of the respondents ones could hinder them from information. This discrepancy of odds might be due knowledge level of respondents toward utilization of iodized salt, study design and study area. This indicates that education has positive relationship for the demand for iodized salt.

From a public health perspective, it is encouraging that almost 90% of the study population already used iodized salt within one year of the introduction of the salt iodization programme(31).Adequate level of iodized salt at household level was significantly increased the odds of proper utilization of iodized salt by 66.8% compared to those households who had inadequate level of iodized salt compared to their counterparts [AOR= 1.668(1.071, 2.597)]. The odds of this was lower than that stud done in Ahfeherom District 3.90 in 2019,which level of iodized salt contents as predictors to the utilization of iodized salt(46).This discrepancy might be due availability of iodized salt at household level and analysis method. This indicates that level of iodized salt has positive relationship for the demand for iodized salt.

It is also important to note that more than 90% of the people knew about the relationship between IDDs and iodized salt(31). Although knowledge on use of iodized salt was very low, which was 30.5% by itself does not guarantee for availability and utilization of adequately iodized salt. Proper utilization of iodized salt were about two and three-folds significantly higher among those who had good knowledge compared to their counterparts [AOR=2.320(1.437, 3.745)].The finding of this study odds higher than odds that studied in Laelay May chew (2.207)in 2015(40).This difference might be due to the fact that the present study period.

In addition to this, the difference in socio-demographic characters and participants might be affecting the proper utilization of iodized salt.The finding of this study odds lower than odds that studied in Mecha(3.8)in 2019,Sidama zone (4.66)in 2016,Dabat Woreda(1.49) in 2017,Asella (4.93)in 2016 and Dega Damot district (5.55)in 2019(14, 38, 41, 45, 53).This discrepancy might be due to information access, study period and study setting. This may show the role of good knowledge on influencing demand and need of properly utilization of iodized salt.

Iodine stability is critical for the success of salt fortification(33).Poor storage of iodized salt also leads to loss of iodine in the salt(54). Iodine content will remain relatively constant if the salt, kept dry, cool and away from light(41).Good practice of iodized salt at household level was significantly increased .The odds of proper utilization of iodized salt by 35.2% compared to those households who had poor practice of iodized salt compared to their counterparts [AOR= 3.352(2.160, 5.202)]. This finding was lower than that study conducted in the Addis Ababa good practice was 76.3%(95%CL72.7,79.8) in 2018(7). This discrepancy might be due to information access and setting area. This indicates that good practice of iodized salt at household level has positive relationship for the demand for iodized salt.

## 5. Conclusion

Proper utilization of iodized salt remains very low, which was 38.4% in the district and does not meet the national goal. Educational status, level of iodized salt, good knowledge and good practice were significantly associated factors with proper utilization of adequately iodized salt in this study.

## 6. Acknowledgment

We would like thank Arsi Universityto give this golden chance. Next, We would like to thank Arsi zone health department for the provision of Iodine test kits. Our appreciation also extends to Hetosa Health Office for providing necessary information and material support. We would like to thank data collectors and supervisors for taking their precious time to collect data. Finally, We would like to thank to participants of the study who have took their time to provide necessary information for my thesis development.

## Authors’ Information

**Abu Tura Buli**^**1**^**\*:**Corresponding Author

**Educational background:** BSc in Environmental health from Haromaya University MPH From Arsi University Health College

**Current Position:**Head of Hetosa Woreda Health office

**Melese Tadesse Aredo** ^**2**^

**Educational background:** BSc in Environmental health from Haromaya University BSc in Clinical Nursing from Arsi University

MPH specialty in Environmental and Occupational health from Addis Ababa University.

**Current Position:** Lecturer at Arsi University College of health sciences

**Hailu Fekadu Demise** ^**2**^:

Educational background: BSc in health officer from Haromaya University General MPH from Addis Ababa University

**Current position:** Assistant professor Research Vice Dean at Arsi University College of health sciences

**Ashenafi Habtamu Regesu**^**2**^:

**Educational background and current position:**

BSc in Environmental Health, From Gondar University

**Current position:** Assistant professor in MPH at Arsi University College of health sciences

## Abbreviations

AIS: Adequate Iodization Salt
AOR: Adjusted Odd Ratio
BSC: Bachelor of Science
COR: Crude Odd Ratio
ICCIDD: International Council for Control of Iodine Deficiency Disorder
IDD: Iodine Deficiency Disorder
IQ: Intelligence Quotient
IS: Iodized Salt
Mr.: Misters
MPH: Master of Public Health
OR: Odd-Ratio
PPM: Parts Per Million
RTK: Rapid Test Kit
SPSS: Statistical Package for the Social Science
UNICEF: United Nation Children Fund’s
USI: Universal Iodization Salt
WHO: World Health Organization

